# *Prochlorococcus* rely on microbial interactions rather than on chlorotic resting stages to survive long-term nutrient starvation

**DOI:** 10.1101/657627

**Authors:** Dalit Roth-Rosenberg, Dikla Aharonovich, Tal Luzzatto-Knaan, Angela Vogts, Luca Zoccarato, Falk Eigemann, Noam Nago, Hans-Peter Grossart, Maren Voss, Daniel Sher

## Abstract

Many microorganisms produce resting cells with very low metabolic activity that allow them to survive phases of prolonged nutrient or energy stress. In cyanobacteria and some eukaryotic phytoplankton, the production of resting stages is accompanied by a loss of photosynthetic pigments, a process termed chlorosis. Here, we show that a chlorosis-like process occurs under multiple stress conditions in axenic laboratory cultures of *Prochlorococcus*, the dominant phytoplankton linage in large regions of the oligotrophic ocean and a global key player in ocean biogeochemical cycles. In *Prochlorococcus* strain MIT9313, chlorotic cells show reduced metabolic activity, measured as C and N uptake by NanoSIMS. However, unlike many other cyanobacteria, chlorotic *Prochlorococcus* cells are not viable and do not re-grow under axenic conditions when transferred to new media. Nevertheless, co-cultures with a heterotrophic bacterium, *Alteromonas macleodii* HOT1A3, allowed *Prochlorococcus* to survive nutrient starvation for months. We propose that reliance on co-occurring heterotrophic bacteria, rather than the ability to survive extended starvation as resting cells, underlies the ecological success of *Prochlorococcus*.

**Importance:** The ability of microorganisms to withstand long periods of nutrient starvation is key to their survival and success under highly fluctuating conditions as is common in nature. Therefore, one would expect this trait to be prevalent among organisms in the nutrient-poor open ocean. Here, we show that this is not the case for *Prochlorococcus*, a globally abundant and ecologically impactful marine cyanobacterium. Instead, *Prochlorococcus* rely on co-occurring heterotrophic bacteria to survive extended phases of nutrient and light starvation. Our results highlight the power of microbial interactions to drive major biogeochemical cycles in the ocean and elsewhere with consequences at the global scale.

## Introduction

Not all microbial cells living in natural environments are equally active. In aquatic environments, up to 90% of the cells do not exhibit measurable metabolic activity (“vitality”), based on dyes (e.g. that assess electron transport) or on uptake assays with labeled substrates (1). Several possible and non-exclusive explanations have been proposed for this heterogeneity. First, “observed differences in activity between cells in natural populations may represent inherent differences in activity between genetically-different organisms, e.g. due to variations in maximum growth rate or the ability to utilize the specific substrate tested. Second, cells might be at different physiological states, e.g. exponentially growing, starved or dying, and thus exhibiting different levels of metabolic activity (2, 3). Third, cells show stochastic fluctuations in their activity, due to noise in gene expression or regulatory networks (4). Finally, some organisms respond to environmental stress by producing resting stages or spores. Such cells often exhibit very low (or undetectable) metabolic activity, yet are viable, namely able to return to an active state and reproduce when environmental conditions return to favorable (5). The presence of such resting stages, together with a fluctuating activity at the single-cell level and the genetic variability found within natural populations, are suggested to promote the survival of the population as a whole (2, 6).

Understanding the factors affecting the metabolic activity (vitality) of phytoplankton is of special interest. These microbial primary producers perform about one-half of the photosynthesis on Earth, providing energy through carbon fixation at the base of the aquatic ecosystem (7). As phytoplankton grow, they take up elements such as nitrogen (N) and phosphorus (P) from the environment, potentially leading to low nutrient concentrations that may constrain the growth of both the phytoplankton themselves and co-occurring organisms (8, 9). Phytoplankton viability, including their ability to survive under conditions of nutrient stress, has been extensively studied, especially for organisms that produce massive blooms that emerge and decline rapidly (reviewed by (10–12)). For example, some bloom-forming cyanobacteria such as *Aphanizomenon* species produce morphologically-distinct spores that show very little photosynthetic activity, yet remain viable in the sediment for long periods of time, providing the inoculum for the next growth season (13). In laboratory cultures of *Synechococcus elegantus* PCC 7942 and *Synechocystis* PCC 6803, two unicellular freshwater cyanobacteria, nitrogen starvation results in a programmed process where cells enter a resting stage, enabling them to survive prolonged periods of stress (14, 15). As part of this process, cells degrade their photosynthetic apparatus in a controlled manner, resulting in a loss of chlorophyll autofluorescence and culture bleaching (a process termed chlorosis). However, the observation that chlorotic cells are viable resting stages is not universal. Chlorotic cultures of *Microcystis aeruginosa* PCC 7806 were shown to contain a small population of non-chlorotic cells with high chlorophyll autofluorescence (described throughout this study as “high-fl”). Only these high-fl cells were suggested to revive after the re-addition of a nitrogen source, while the low-fl cells are presumably dead (16). Chlorotic cells were also observed in eukaryotic phytoplankton, however it is not yet clear to what extent such cells remain viable, as it may depend on the specific organism and stress conditions (11, 17, 18).

*Prochlorococcus* is a pico-cyanobacterium that is extremely abundant in the oligotrophic oceans, performing an estimated ~8.5% of global ocean photosynthesis (19). The carbon fixed by *Prochlorococcus*, which are estimated to produce up to 75% of the daily photosynthetic carbon in the surface layer of the Pacific subtropical gyre (20), can then be utilized by co-occurring heterotrophic bacteria. *Prochlorococcus* cells in the oceans exhibit extremely high genetic diversity (21), and some of this diversity has been linked with their ability to grow under conditions of extreme nutrient limitation (e.g. (22, 23)). It has therefore been suggested that this genetic diversity enables *Prochlorococcus* as a group to thrive across a wide variety of oceanic conditions (24). While the physiological and transcriptional responses of multiple *Prochlorococcus* lineages to short-term nutrient starvation have been extensively studied (e.g. (22, 25–29)), little is known about their ability to survive more than a few days under such conditions. A study on the response of *Prochlorococcus* strains to a different type of stress, extended darkness (i.e. C starvation), has shown that these organisms can survive light starvation only for a limited time (30). In these experiments, low-fl cell populations reminiscent of chlorotic cells in other cyanobacteria appeared after the light-starved cultures were re-exposed to light (30). Therefore, phenotypic evidence exists that *Prochlorococcus* can undergo a chlorosis-like process, yet whether these chlorotic cells are active, and whether they are resting stages that can resume growth when conditions are favorable, is currently unknown. Our experiments were therefore designed to answer the following questions: i) Do *Prochlorococcus* respond to long-term nutrient starvation by producing chlorotic cells? ii) If so, are such cells metabolically active (vital) and are they able to reproduce and grow when stress conditions end (viable)? To address these questions, we used fluorescence-activated cell sorting (FACS) to obtain distinct chlorotic sub-populations from axenic and unialgal laboratory cultures of *Prochlorococcus* which were pre-incubated with isotopically-labelled tracers for photosynthesis (H^13^CO_3_) and nutrient uptake (^15^NH_4_^+^) and we visualized their activity using Nanoscale Secondary Ion Mass Spectrometry (NanoSIMS). This method enabled us to measure photosynthesis and N uptake at a single cell resolution by quantifying the change in isotopic ratios (31, 32). Our results show that while *Prochlorococcus* do undergo a chlorosis-like process, with some of the chlorotic cells still photosynthesizing and taking up NH_4_^+^, the chlorotic cells are unable to re-grow and thus do not represent resting stages. Instead, co-culture with heterotrophic bacteria enables *Prochlorococcus* to survive long-term stress even without producing resting stages.

## Results and discussion

### Emergence of chlorotic sub-populations in *Prochlorococcus* cultures

As *Prochlorococcus* batch cultures reach stationary stage and start declining, the green color of the cultures disappears, and sub-populations of cells emerge with lower chlorophyll autofluorescence that can be identified by flow cytometry (Fig. 1A, B). This phenomenon is observed in strains from all major cultured ecotypes, as well as in a marine *Synechococcus*, strain WH8102 (Fig. 1C). In the high-light adapted strain *Prochlorococcus* MIT9312, lower chlorophyll populations emerged in batch cultures that reached stationary stage due to both N and P limitation, although the timing of sub-population emergence and the forward light scatter and chlorophyll autofluorescence (analyzed by flow cytometry) were different under the two nutrient stresses (Fig. S1A, B, (33)). Cells with lower chlorophyll autofluorescence also appeared in populations of another strain, the low-light adapted MIT9313, when these cultures were inhibited in a co-culture with high cell densities of the heterotrophic bacterium *Alteromonas* HOT1A3 (Fig. S1C, D (34)). Thus, the emergence of populations of cells with lower chlorophyll autofluorescence under a variety of stress conditions is a pervasive phenomenon across marine pico-cyanobacteria. We focused our experiments aiming to better characterize this phenomenon on *Prochlorococcus* strain MIT9313, as the response to stress in this strain has been extensively studied (e.g. (22, 26, 27, 34–36)). Additionally, in this strain, three clearly separate sub-populations can be observed when cultured in Pro99 media, facilitating the sorting and subsequent NanoSIMS analyses (Fig. 1B, referred to throughout the study as high-, mid- and low-fl populations).

**FIG 1.**
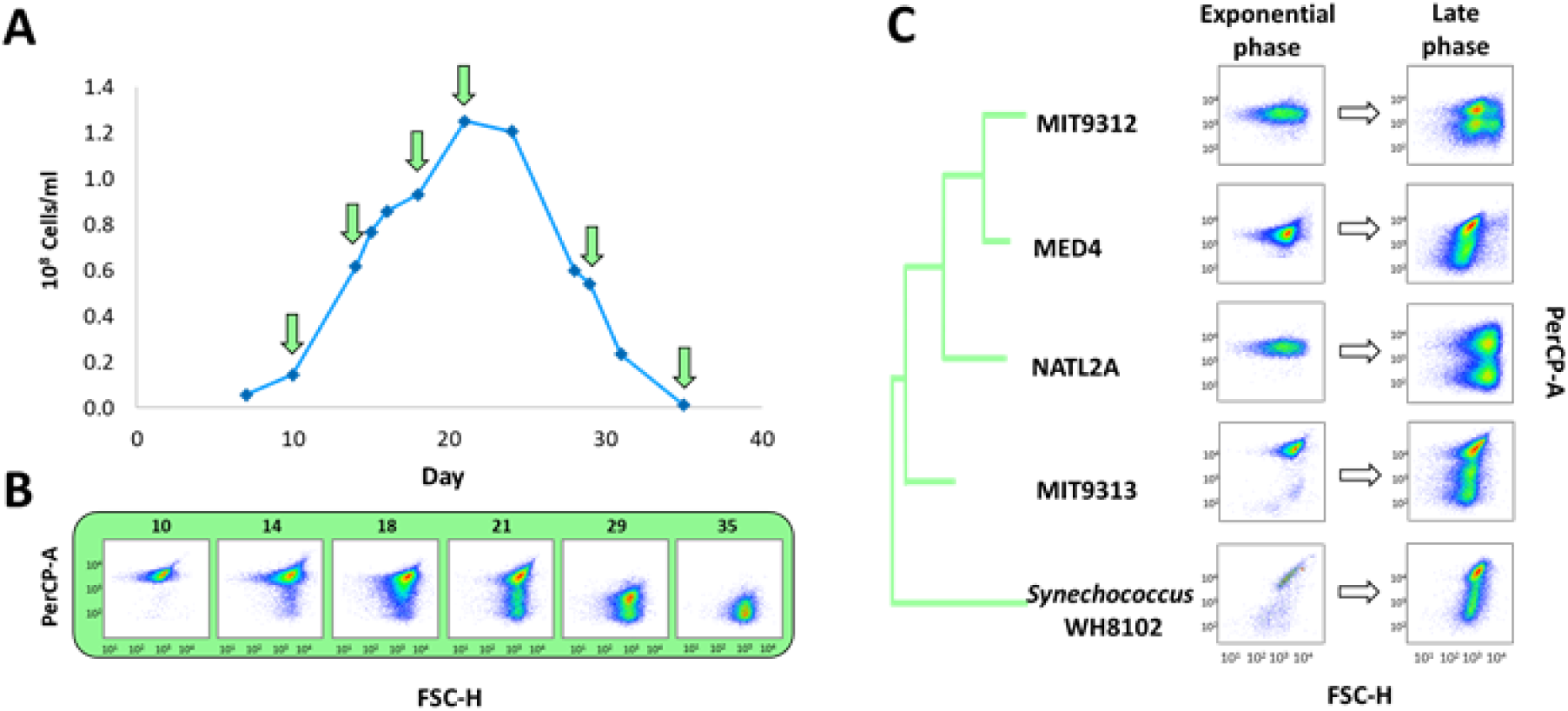
Emergence of chlorotic sub-populations in *Prochlorococcus* batch cultures as measured by flow cytometry. (A) A representative growth curve of an axenic culture of MIT9313, grown in Pro99. The arrows mark the days shown in panel B. (B) Flow cytometry scattergrams at the marked time-points from the MIT9313 culture. The x-axis is Forward Scatter (FSC, a proxy for cell size) and the y-axis is the chlorophyll autofluorescence of the cells (Per-CP). The emergence of chlorotic sub-population observed from the late exponential phase (Day 18). (C) Chlorotic sub-population observed in ageing batch cultures of *Prochlorococcus*, belonging to different ecotypes: High-Light adapted MED4 (HLI), MIT9312 (HLII) and Low-Light adapted NATL2A (LLI) and MIT9313 (LLIV). In all strains, the chlorotic G cells begin to emerge at late growth stage, becoming dominant in declining cultures, while in the exponential phase only one population can be observed. Additional growth curves for this strain and for others, including replicates and standard deviations, as shown in Fig. 3 and 4 and in Fig. S1-3, and S5-6.

### Assessing the metabolic activity of sorted chlorotic sub-populations

We next asked whether the high, mid- and low-fl populations differ in their vitality, measured here as their photosynthesis and nutrient uptake rates (incorporation of H^13^CO_3_^-^ and ^15^NH_4_^+^, respectively). The uptake ratio of labeled versus unlabeled nutrients were then used to calculate the metabolic activity of the sorted cells (Table 1). As shown in Fig. 2 and Table 1, the mean uptake of both H^13^CO_3_^-^ and ^15^NH_4_^+^ was highest in the high-fl population, followed by the mid and low-fl populations, with the latter population indistinguishable from the control, i.e. glutaraldehyde-killed cells. We have repeated the entire workflow in an independent experiment, and the results are very similar (Fig. S2A, B, Table 1). These results are reminiscent of observations in several eukaryotic phytoplankton (18).

**Table 1.**
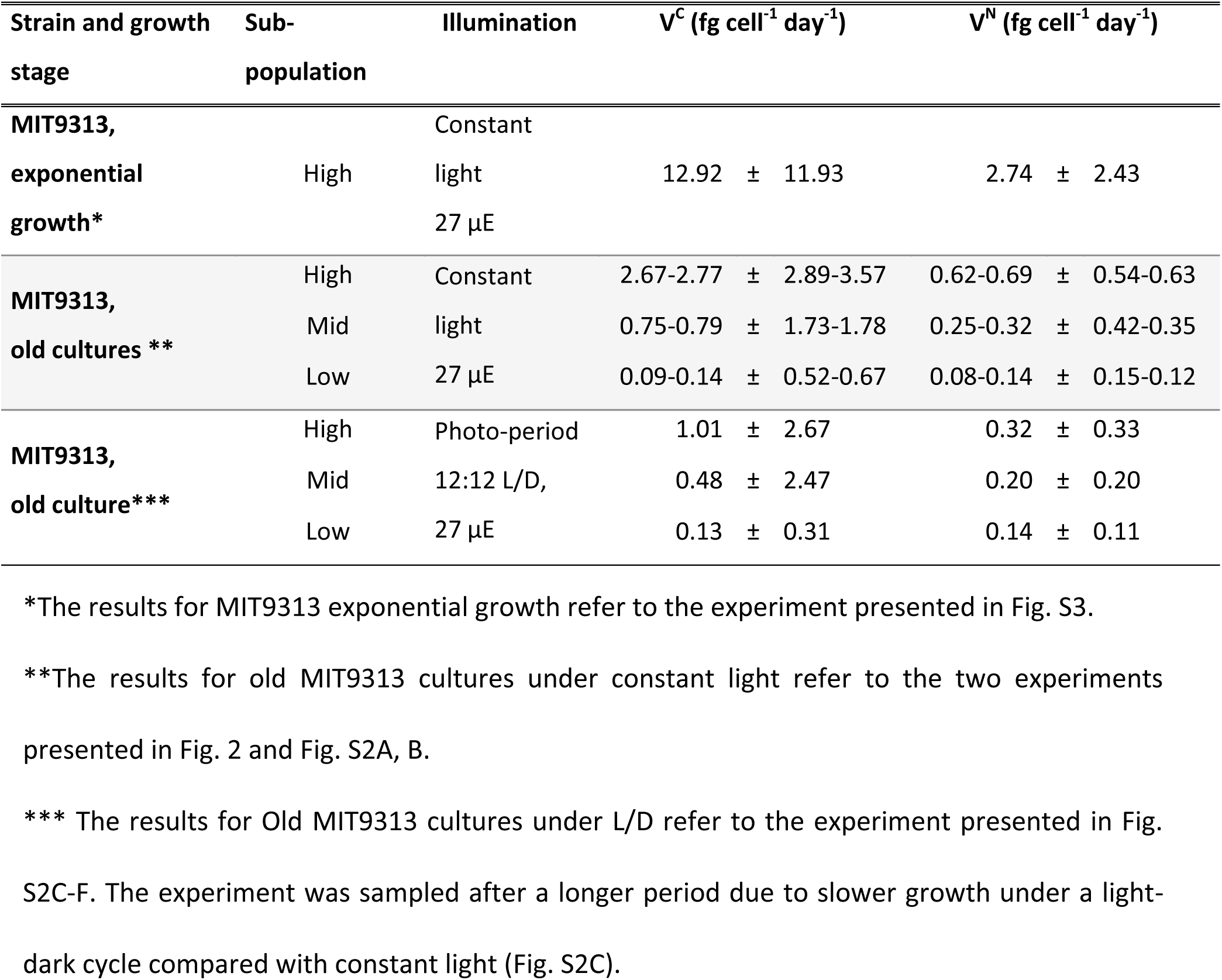
Calculated mean C and N uptake rates from the experiments performed with *Prochlorococcus* MIT9313. The means and standard deviation were calculated from the uptake rates values of single cells in each experiment.

**FIG 2.**
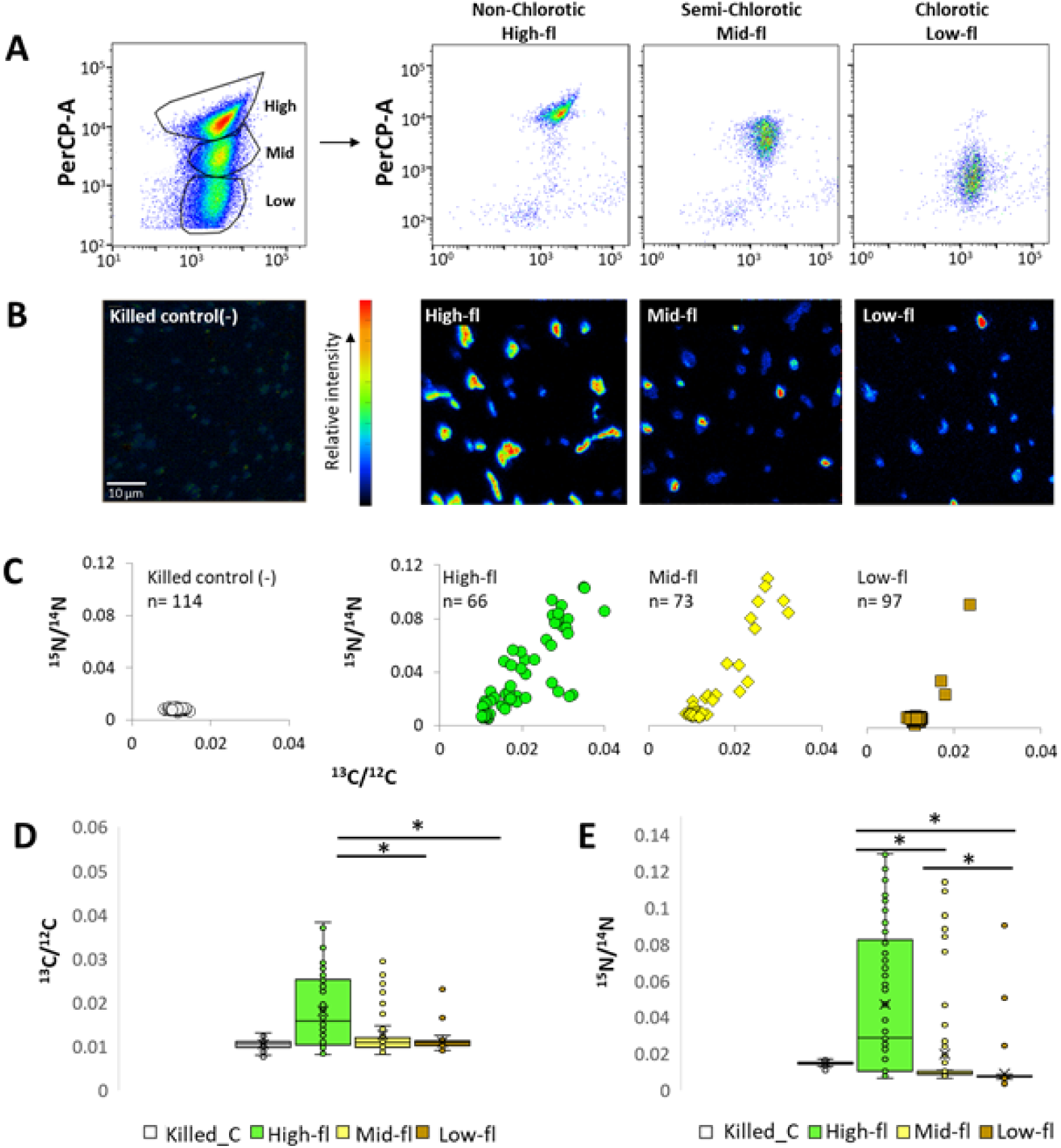
Metabolic activity of sorted sub-populations by NanoSIMS (A) Flow cytometry scatterplots before and after sorting of three distinct sub-populations (high, mid and low-fl) of an aging *Prochlorococcus* MIT9313 culture, detected by flow cytometry. The cultures were grown for 30 days in Pro99 and labeled with H^13^CO_3_^-^ and ^15^NH_4_^+^ for 18h. (B) NanoSIMS images of ^15^N/0^12^C analysis of killed cells (negative control) and high, mid and low-fl cells after sorting. (C) Scatterplot of ^13^C/^12^C and ^15^N/^14^N ratios obtained from NanoSIMS analysis of each sub-population. (D, E) Boxplots of the ^13^C/^12^C and ^15^N/^14^N enrichment in each sub-population. Lines represent the median, X represents the mean, box borders are 1^st^ quartiles and whiskers represent the full range.

The mean uptake rates for glutaraldehyde killed cells (control) were 0.06±0.15 fg cell-1 day-1 for C and 0.18±0.02 fg cell-1 day-1 for N, and most likely depict the absorption of the label by non-specific binding or diffusion.

Within each of the populations, cell-cell heterogeneity was observed in both ^13^C and ^15^N uptake (Fig. 2, Fig. S2A,B). Within all of the populations (including the high-fl), some cells were inactive, and this could not be explained by the limited purity of the FACS-sorting procedure (Table S1, supplementary text). The coefficients of variation in C and N uptake rates were within the range shown for other organisms, or higher (Table S2 (32, 37)). Similar levels of heterogeneity (primarily in N uptake) were also seen in cells grown under a 12:12 light-dark cycle, where the *Prochlorococcus* cell-cycle follows a diel rhythm, suggesting that this heterogeneity is not due to different stages of the cell cycle or the diel cycle (Table S2, Fig. S2C-F). Cell-cell heterogeneity was also observed in cells from an exponentially-growing, nutrient-replete culture (Fig. S3, Table S2), suggesting that this heterogeneity is not exclusively limited to ageing or stressed cells. This is in accordance with studies assessing the vitality of *Prochlorococcus* cells using various dyes, which consistently show that a significant fraction of the cells in laboratory cultures are inactive or potentially dead (38, 39).

In addition to differing in their chlorophyll autofluorescence and metabolic activity, the high-, mid- and low-fl cell populations also differ by their forward and side light scatter properties, which are related to cell size and (in larger cells) morphological complexity (Fig. S4A, B). In agreement with these observations, cells sorted from the high-fl population and observed by SEM (Scanning Electron Microscopy) were 20-30% larger than those from the mid- and low-fl populations (Fig. S4C-E).

### Evaluating the viability of sub-populations

We next asked whether the low-fl cells are viable resting stages. We tested this indirectly by determining the ability of *Prochlorococcus* MIT9313 cells, cultured in Pro99 media, to grow upon transfer to new growth media at different times during exponential growth and upon culture decline. As shown in Fig. 3A, only cells from cultures where the high-fl cells were dominant could grow when transferred to new growth media. No growth was observed upon transfer of cells from stationary or declining cultures where no high-fl cells were observed. Intriguingly, the presence of high-fl cells was not enough to ensure culture growth (e.g. day 34 in Fig. 3A). This is consistent with a previous study showing that cells belonging to a different *Prochlorococcus* strain, MED4, that were incubated for three days in the dark, were unable to resume growth after return to light despite showing no clear difference in the chlorophyll autofluorescence (30). The probability of growth after transfer did not depend on the number of transferred cells (40), with as many as 2.5×10^7^ cells/ml failing to grow after transfer during culture decline (cells at ~1/10 of this density grew after being transferred during exponential stage). Thus, non-chlorotic cells (defined as being within the range of chlorophyll autofluorescence exhibited by exponentially-growing cells) are not necessarily viable.

**FIG 3.**
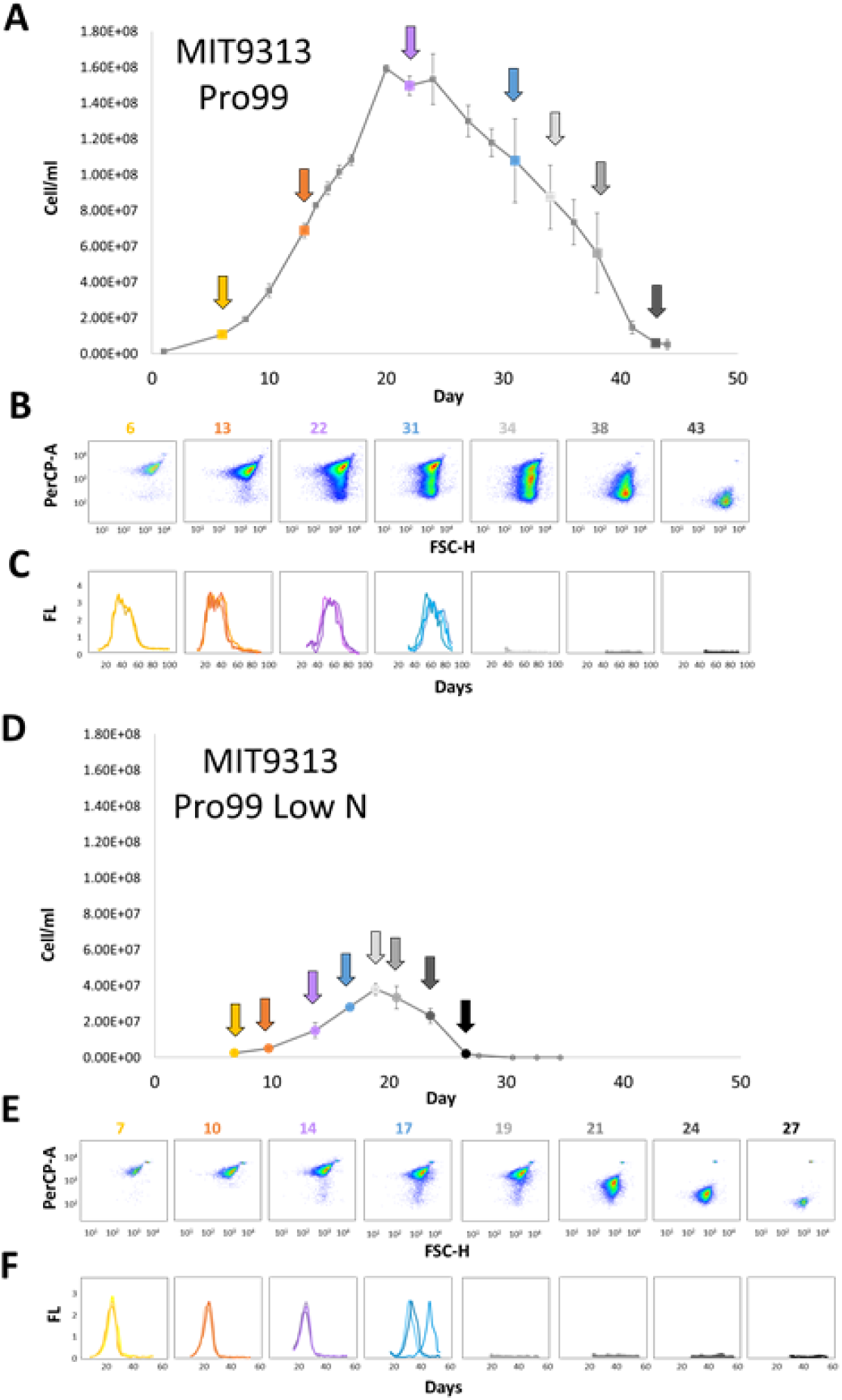
Time-dependent changes in viability of cells transferred into fresh media at different life cycle stages of a batch culture for MIT9313 in Pro99 (A-C) and when stationary stage is induced by N starvation (D-F). (A, D) Growth curves of a MIT9313 culture, in Pro99 media commonly used for *Prochlorococcus* culturing (A) and under conditions where stationary stage is induced by nitrogen starvation (panel (D), 2:1 N/P ratio in the growth media, (42)). Colored circles indicate the times point at which triplicate 1ml samples were transferred into 20 ml fresh media. (B, E) Flow cytometry scatterplots of the culture shown in panels (A) and (D). Note that, under conditions of N starvation, the cultures shift rapidly from being comprised primarily of high-fl cells (day 19 in panel (E), early stationary phase) to mainly mid-fl cells, with essentially no high-fl cells (day 21 in panel (E). (C, F) Growth curves of cells being transferred at different times to new, nutrient-replete media (assessed via bulk culture fluorescence). In these plots, each line shows a replicate culture. Cells could not re-grow when transferred after more than 31 days in Pro99 and 17 days of nitrogen starvation. This suggests that, in this strain, high-fl cells are not necessarily viable.

One problem with performing experiments in Pro99, which is commonly used to culture *Prochlorococcus*, is that the conditions causing cells to reach stationary stage are not always clear (e.g. (41). We therefore repeated these experiments under conditions where entry into stationary phase is induced by N or P starvation (Fig. 3B, Fig. S5, (42)). When entry into stationary stage was induced by N or P starvation, chlorotic cells appeared much faster, and the cultures became non-viable much earlier – essentially immediately after the cessation of exponential growth. Similar results were obtained with a different strain of *Prochlorococcus*, MIT9312 (Fig. S5A, B). However, a marine *Synechococcus* (strain WH8102) behaved differently, surviving N starvation much longer and being able to re-grow in nutrient replete media long after the culture started declining, and when essentially all cells were chlorotic (Fig. S5D). This is reminiscent of the ability of (presumably axenic) cultures of two freshwater cyanobacteria, *Synechococcus* PCC 7942 and *Synechocystis* PCC 6803, to revive after extended N starvation (14, 15).

The inability of axenic *Prochlorococcus* strains to survive long-term nutrient starvation was surprising, and we therefore hypothesized that their survival would be enhanced by interactions with co-occurring heterotrophic bacteria. Indeed, when co-cultured with a heterotrophic bacterium, *Alteromonas* HOT1A3 (34, 43), *Prochlorococcus* strains representing all major cultured ecotypes were able to re-grow after 60 days of N and P stress, whereas all axenic strains failed to do so (Fig. 4, Fig. S6A, B). Interestingly, strain MIT9313, which was initially inhibited by this *Alteromonas* strain (Fig. 4A, (34, 44)), was also able to survive long-term starvation in co-culture, suggesting that fundamentally different interactions occur during exponential growth compared to long-term, presumably nutrient-limited growth. These results are consistent with the ability of heterotrophic bacteria to extend the survival time of different *Prochlorococcus* strains under conditions of constant darkness (albeit for only several days, (30)) and with the ability of different heterotrophic bacteria to support the long-term viability of batch cultures of *Synechococcus* WH7803 (45). We observed similar results when *Prochlorococcus* cultures were transferred from nutrient-replete media into sterile sea water with no added nutrients (Fig. S6C, D). This suggests that the ability of the heterotrophs to support *Prochlorococcu*s survival is due to nutrient remineralization rather than to the detoxification of potential waste products, although a potential role for de-toxification cannot be ruled out (45).

**FIG 4.**
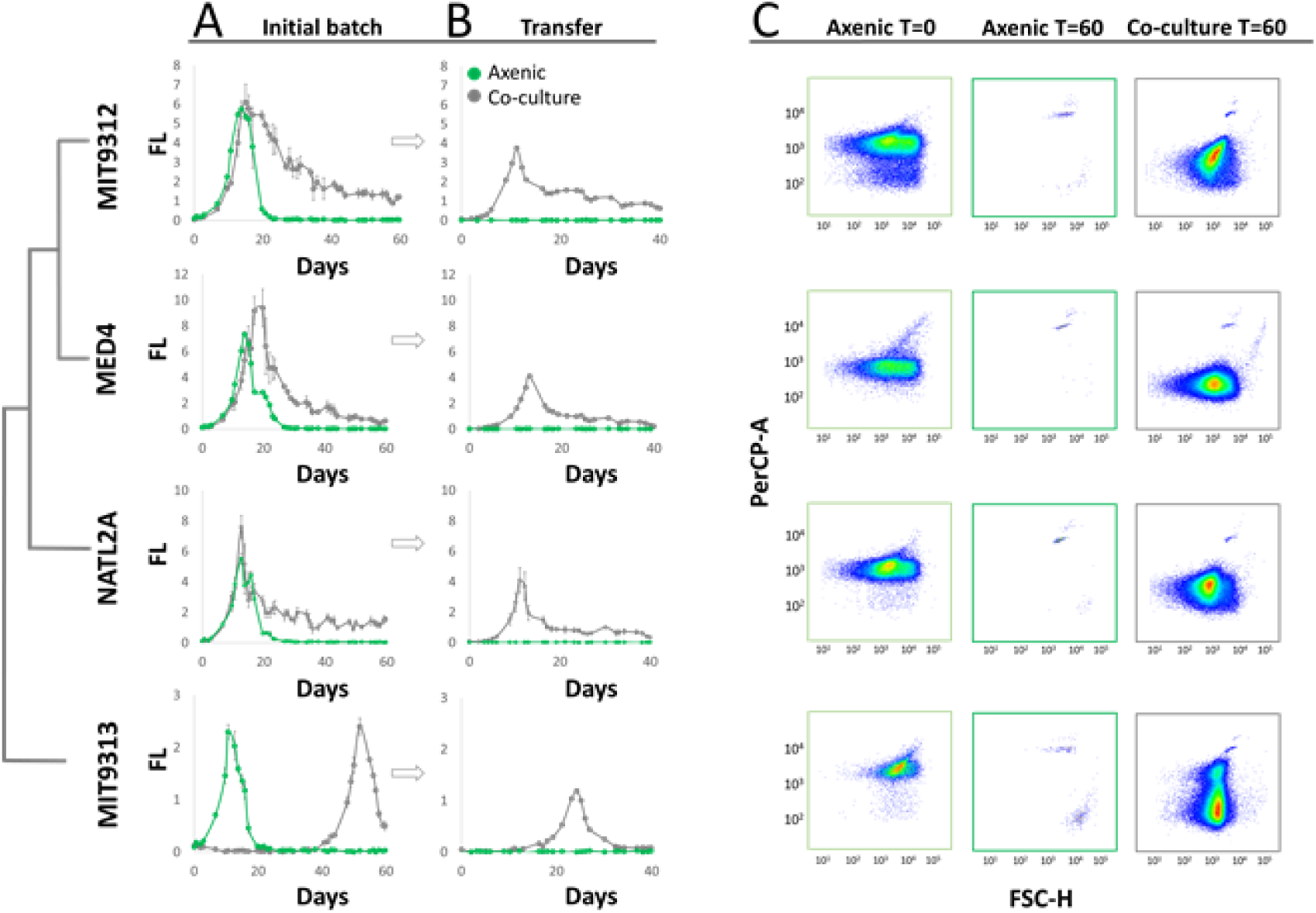
Co-culture with a heterotrophic bacterium, *Alteromonas* HOT1A3, enables multiple *Prochlorococcus* strains to survive long-term N starvation. (A) 10^6^ axenic *Prochlorococcus* cells ml^-1^ from different strains were incubated alone (green line) or with the addition of 10^7^ *Alteromonas* HOT1A3 cells ml^-1^ in low-N media (grey line). Bulk culture fluorescence was recorded as a proxy for cell growth, and 1ml from each culture was transferred into 25ml fresh Pro99 media after 60 days. Panel (B) The transferred cultures were recorded for additional 40 days. Error bars are standard deviation from triplicate cultures. The late growth of MIT9313 in co-culture is the “delayed growth” phenotype described in (34, 44). (C) Flow cytometry scattergrams of the cultures shown in panel A. Co-culture with *Alteromonas* increases the number of *Prochlorococcus* cells seen by flow cytometry, suggesting a reduction in the magnitude of mortality (cell lysis). A decrease in per-cell chlorophyll autofluorescence is still seen, suggesting co-culture does not completely inhibit the process of chlorophyll degradation.

### Stress survival in pico-cyanobacteria: why is *Prochlorococcus* different?

In this study, we demonstrate that phenotypic heterogeneity between clonal *Prochlorococcus* cells occurs at multiple “scales”. In exponentially growing axenic laboratory cultures of two strains, MIT9313 and MED4, C and N uptake rates differ significantly between individual cells (summarized in Table S2). This variation is independent of genetic variability. Additionally, as axenic cultures become stressed, a larger phenotypic change occurs as cells lose their chlorophyll auto-fluorescence and become chlorotic. Under these experimental conditions, most cells are inactive (primarily in the low-fl population, as measured in strain MIT9313), although we cannot rule out that even low-fl cells still retain a residual level of activity not detectable by the nanoSIMS. Nevertheless, some cells from the chlorotic populations retain at least part of their photosynthetic capacity, and indeed can fix carbon and take up NH_4_. Yet, in our experiments, they do not re-grow when condition become more favorable. In *Synechococcus elegantus* PCC 7942, chlorotic cultures retain approximately 0.01% of their photosynthetic activity, as well as a residual level of protein translation, although it remains unclear whether this is a process shared by all cells in the culture or whether this activity is only due to a small subset of more active cells (14). The clear difference between the ability of axenic *Synechococcus elegantus* PCC 7942 and *Synechocystis* PCC6803 to survive long-term N starvation, and the inability of axenic *Prochlorococcus* cultures to do so, suggests an inherent difference in the physiology and genomic functional capacity between these unicellular cyanobacteria.

Entry into chlorosis in *Synechocystis* is a regulated process that involves the organized degradation of the phycobilisomes in parallel with an increase in the storage products glycogen and polyhydroxybutyrate (PHB) (15). The photosynthesis apparatus of *Prochlorococcus* is different from that of other cyanobacteria, using unique chlorophyll a2/b2 binding proteins rather than phycobilisomes (46). Indeed, *Prochlorococcus* lack orthologs of the nblA gene required for phycobilisome degradation during chlorosis (15). Additionally, while *Prochlorococcus* likely use glycogen as a C storage pool (47), they lack the phaA-C and phaE genes required for PHB biosynthesis and which are induced in *Synechocystis* PCC 6803 under chlorosis (although these genes are not required for revival from chlorosis (15)). Taken together, these differences suggest that *Prochlorococcus* lack the genetic toolkit employed by *Synechocystis* PCC6803 and *Synechococcus elegantus* PCC7942 to enter into a resting stage. Thus, chlorotic cells in *Prochlorococcus* are not resting stages. However, we note that *Synechococcus* WH8102 also lacks the gene nblA gene, yet can survive N starvation much longer in axenic culture than the tested *Prochlorococcus* strains can (Fig. S5D). This suggests that additional molecular mechanisms besides nblA-mediated degradation of the phycobilisomes are important for long-term survival of nutrient starvation in *Synecococcus* WH8102.

If *Prochlorococcus* are indeed incapable of producing resting stages in response to nutrient or light starvation, what are the evolutionary drivers of this phenotype, and what are the consequences for the dynamics of *Prochlorococcus* populations in the ocean? While the open oligotrophic ocean is often considered a relatively stable environment, nutrient concentrations do fluctuate (8), and phytoplankton (including *Prochlorococcus*) inhabiting these waters show multiple signs of nutrient stress (48). Many of the microbes that live in such environments comprising a large fraction of the surface ocean have small, highly streamlined genomes (49) and this has been suggested to be an adaptation to low nutrient concentrations (24, 49, 50). Therefore, it is possible that the lack of resting stages is a result of this genome streamlining - the genomes of *Synechococcus elegantus* PCC7942 and *Synechocystis* PCC6803 are ~3.2mbp and ~4 mbp with their plasmids, respectively, compared to ~1.4-2.5 mbp for *Prochlorococcus* strains.

### Surviving nutrient stress “with a little help from my friends”

The ability of *Prochlorococcus* to thrive under conditions of extreme nutrient limitation is often explained by their small cell size (increasing their biomass-specific diffusion), their generally low nutrient requirements, and their specific metabolic strategies to minimize the per-cell elemental quotas (51–53). However, these mechanisms appear not to work in axenic laboratory cultures, and thus we propose that interactions with co-occurring microorganisms enable *Prochlorococcus* to survive when nutrient-saving mechanisms are not sufficient, as suggested for its close relative *Synechococcus* (45). This may take the form of recycling of inorganic nutrients by the heterotrophic bacteria, as well as possibly by the production of organic compounds that contain elements such as N or P. Indeed, *Prochlorococcus* can compete with heterotrophic bacteria for amino acids (54). Importantly, the utilization of organic compounds (mixotrophy) may provide *Prochlorococcus* also with carbon, sulfur or energy sources, and may potentially help them survive also light starvation (e.g. (30, 55–59)).

Regardless of the specific forms of dissolved organic matter being utilized by the cells, and on the exact mechanism enabling the cells to survive long-term nutrient starvation in co-culture, the lack of any observed mechanism for the production of resting stages by *Prochlorococcus* may be considered another manifestation of the “Black Queen Hypothesis”. This hypothesis states that microorganisms “outsource” essential survival mechanisms such as detoxification of reactive oxygen species to the surrounding microbial community (60). These forms of microbial interactions likely affect the distribution and activity of *Prochlorococcus* on a global scale. The increased survival of *Prochlorococcus* under harsh conditions, supported by its associated heterotrophic bacteria, may enable it to remain active at the single cell level even during long periods of unfavorable conditions (61, 62). Thus, the tight interactions between *Prochlorococcus* and its bacterial “supporters” likely affects photosynthesis and carbon cycling at the base of the aquatic foodweb, with potentially profound implications for overall oceanic productivity and carbon cycling.

## Material and Methods

### *Prochlorococcus* growth and Stable Isotope Incubations

Axenic *Prochlorococcus* strains were grown in Pro99 media under constant cold while light (27 μE) at 22 °C. Axenicity of the strains was tested routinely, as well as before all major experiments, using test media (ProMM, (40)). Additionally, no evidence for contaminating heteroteophic cells was observed in flow cytometry, scanning electron microscopy or when axenic *Prochlorococcus* cultures were used as negative controls for 16S amplicon sequencing. Bulk chlorophyll fluorescence (FL) (ex440; em680) was measured almost daily using a Fluorescence Spectrophotometer (Cary Eclipse, Varian). In parallel, samples for flow cytometry were taken for cell numbers. When three distinct sub-populations appeared in the flow cytometry, the cultures were labeled with 1mM Sodium bicarbonate-^13^C and 1mM Ammonium-15N chloride (Sigma-Aldrich, USA) for 18-24 hours. The optimal incubation time based on preliminary isotope labeling experiments with *Prochlorococcus* MED4, showing that uptake is identified already after 3 hours and is linear until 24 hours under our growth conditions (Fig. S7). Incubations were stopped by fixing 2 ml of the culture with 2X EM grade glutaraldehyde (2.5% final concentration, Sigma) and subsequently storing at 4 °C until the sorting analysis. Non-labeled cells that were killed before labeling (by adding 2.5% glutaraldehyde) were used as a negative control.

### Cell Sorting and Filtration

Sorting of sub-population was carried out using a BD FACSAria III sorter (BD Biosciences) at the Life Sciences and Engineering Infrastructure Center, Technion, Israel. Each sample was sorted for 3 sub-populations: Non-chlorotic (High-fl), Semi chlorotic (Mid-fl) and Chlorotic (Low-fl) (Fig. 2A). The sorting gates for each sub-population were determined from the population observed in forward scatter (FSC, a proxy for cell size) and auto-fluorescence (PerCP, chlorophyll auto-fluorescence). After sorting, the sorted sup-population were gently filtered on 13 mm diameter polycarbonate filters (GTTP, 0.2 μM pore size, Millipore, MA), washed twice with sterile sea water and air-dried. The filters were cut into two parts. One half was stored at 4°C until nanoSIMS analyses and the other half is for Scanning Electron Microscopy.

### Nanoscale secondary ion mass spectrometry (nanoSIMS) and data analysis

The samples were coated with a layer of ca. 30 nm gold with a Cressington 108 auto sputter coater (Watford, United Kingdom). Random spots were employed for NanoSIMS analyses. SIMS imaging was performed using a NanoSIMS 50L instrument (Cameca, Paris, France) at the Leibniz-Institute for Baltic Sea Research Warnemünde (IOW). A ^133^Cs^+^ primary ion beam was used to erode and ionize atoms of the sample. Images of secondary electrons, ^12^C^-^, ^13^C^-^, ^12^C^14^N^-^ and ^12^C^15^N^-^ were recorded simultaneously using mass detectors equipped with electron multipliers (Hamamatsu). The mass resolving power was adjusted to be sufficient to suppress interferences at all masses allowing, e.g. the separation of ^13^C^-^ from interfering ions such as ^12^C^1^H^-^. Prior to the analysis, sample areas of 50×50 µm were sputtered for 2 min with 600 pA to erode the gold, clean the surface and reach the steady state of secondary ion formation. The primary ion beam current during the analysis was 1 pA; the scanning parameters were 512×512 pixels for areas of 30x30 to 48x48 µm, with a dwell time of 250 µs per pixel. 60 planes were analysed.

### Analyses of NanoSIMS measurements

All NanoSIMS measurements were analysed with the Matlab based program look@nanosims (63). Briefly, the 60 measured planes were checked for inconsistencies and all usable planes accumulated, regions of interest (ROI’s) (i.e. *Prochlorococcus* cells and filter regions without organic material for background measurements) defined based on ^12^C^14^N mass pictures, and ^13^C/^12^C as well as ^12^C^15^N/^12^C^14^N ratios calculated from the ion signals for each region of interest.

### Uptake rate calculation

Uptake rate was estimated using the following equation, based on that of (64), as follows:

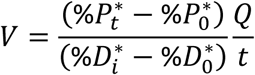

Where *%P*_t_* is the concentration (atom %) of the heavy isotope in the particulate matter at the end of the incubation, *%D*_i_* is the concentration of the dissolved tracer added to the incubation (and assumed not to change over the short incubation time), and *%P*_0_* and *%D*_0_* are the natural heavy isotope concentrations in the particulate and dissolved matter, respectively. We estimated *Q*, the cell quota (in fg cell^-1^) of C or N, based on measurements of the biomass of MED4 and MIT9313 (66 fg cell^-1^and 158 fg cell^-1^, respectively, (65)) and assuming that C comprises 50% and N comprises 7.5% of the cell biomass. For heavy isotopes concentration in the particulate and dissolved phases before incubation we used the natural values for isotopic ratios of ^13^C and ^15^N (1.12% and 0.37% respectively). For the experiment shown in Fig. S7, we measured the NH_4_^+^ concentration in the media and added the ^15^N tracer to 50% final concentration. Since all other experiments were performed in declining cultures we assumed that the NH_4_^+^ was depleted from the media, and thus *%D\**_*t*_ was defined as 90%, based on previous measurements of NH_4_^+^ concentrations in old cultures. We used a value of 50% for the initial percentage of ^13^C, based on dissolved inorganic carbon (DIC) measurements (42). For the terminal concentrations of ^15^N and ^13^C in the particulate phase (*%P^*^_t_*) we used the values of ^13^C/^12^C and ^15^N/^14^N that were obtained from the NanoSims analysis of the cells. ^13^C/^12^C and ^15^N/^14^N below the natural values resulted with negative uptake values, and were treated as zero uptake.

Mean and standard deviation of C and N uptake rates were calculated from the uptake rate values of individual cells (Table 1). The uptake rate values were not corrected for negative control (killed cells), which are presented for comparison in table 1. Since ^13^C/^12^C and ^15^N/^14^N values of individual cells were not normally distributed, for significance analysis we used non-parametric tests (Mann-Whitney and Kruskal-Wallis tests) performed using the Real Statistics Resource Pack software (Release 5.4 www.real-statistics.com).

### Scanning Electron Microscopy

Immediately after filtration, filters dehydrated in an ethanol series of 30%, 50%, 70%, 80%, 90% and 100% (vol/vol) ethanol (dilutions were in deionized water) for 10 min each. Samples were then dried, mounted on stubs with carbon tape and coated with 5 nm with gold. Cells were obtained on a ZEISS SigmaTM SEM, by using a SE2 detector (2–2.5 kV, WD = 8 mm).

## Acknowledgements

We thank Annett Grüttmüller for NanoSIMS routine operation. This study was supported by grant RGP0020/2016 from the Human Frontiers Science Program (to MV, HPG and DS) and by grant number 1635070/2016532 from the NSF-BSF program in Oceanography (NSFOCE-BSF, to DS). The NanoSIMS at the Leibnitz-Institute for Baltic Sea research in Warnemuende (IOW) was funded by the German Federal Ministry of Education and Research (BMBF), grant identifier 03F0626A.

## Author contributions

DRR, DA, TLK, AV, MV and DS designed experiments, DRR, DA, TLK, LZ, NN and DS performed experiments and field analyses, DRR, DA, TLK, AV, and FE performed NanoSIMS analyses, DRR, DA, TLK, AV, LZ, FE, NN, HPG, MV and DS analyzed results, DRR, DA, TLK and DS wrote manuscript with contributions from all authors.

## Competing interests

The authors declare no competing interests

## Materials and Correspondence

Please send requests for materials or other correspondence to Daniel Sher

## Notes

### Competing Interest Statement

The authors have declared no competing interest.

### Summary of Updates

The manuscript was shortened significantly, and the part dealing with field observations has been removed for clarity (it will form the basis of a separate manuscript)

