## Supporting Information for "*Prochlorococcus* rely on microbial interactions rather than on chlorotic resting stages to survive long-term nutrient starvation"

Running title: Interactions and starvation in *Prochlorococcus*

*Dalit Roth-Rosenberg, Dikla Aharonovich, Tal Luzzatto-Knaan contributed equally to this study.

**Supplementary text**

**Can mis-sorted cells explain the presence of non-active cells in the high-fl population, and of active cells in the low- and mid-fl populations?**

FACS-sorting high-, mid- and low-fl populations from *Prochlorococcus* cultures resulted in samples that are highly enriched in the sorted populations. However, the sorted populations are not completely pure, with some incorrectly sorted cells observed when the sorted populations were re-analyzed by flow cytometry (e.g. mid- or low-fl cells in the sorted high-fl population, Table S1A). These incorrectly sorted cells could affect the interpretation of the single-cell uptake rates, for example, the presence of inactive cells in the high-fl population could be interpreted either as a real biological phenomenon (cells with high autofluorescence that are nevertheless inactive) or as the result of incorrectly sorted cells belonging to the mid- or low-fl populations (assuming the mid- and low-fl cells are inactive). To test this hypothesis, we first determined, for each cell in the NanoSIMS analysis, whether it was active or inactive, defining inactive cells as those with C and N uptake rates in the range of the control, i.e. glutaraldehyde killed cells (Table S1b, Killed cells C_max_ = 0.765 fg cell^-1^ day^-1^ and N_max_ = 0.215 fg cell^-1^ day^-1^. We then compared the observed number of active cells to the expected number of active cells, based on the number of high-, mid- and low-fl cells after FACS sorting, and assuming only the high-fl cells are active. For the datasets shown in Fig. 2 and Fig. S2B, the hypothesis that the number of active cells could be explained by the number of high-fl cells after sorting was rejected (X^2^ test, DF=2, p<0.01, X^2^=24.4 and n=236 cells for Fig 2, X^2^=336.1 and n=415 cells for Fig. S2). There were fewer active cells in the high-fl populations than expected (68% and 64% active cells Table S1b, compared to 94% and 84.5% high-fl cells, respectively, in the high-fl subpopulation Table S1A), suggesting that some high-fl cells are inactive. Conversely, there were more active cells than expected in the mid- and low-fl population (Table S1), suggesting that some cells in these populations can be active.

**Supplementary Table S1A: Cell counts and purity of the sorted cells**

|  |  | Sorted sub-populations  *MIT9313 old | | | Sorted sub-populations  **MIT9313 old | | |
| --- | --- | --- | --- | --- | --- | --- | --- |
| **% Purity and number of sorted cells** |  | **High** | **Mid** | **Low** | **High** | **Mid** | **Low** |
|  | **High** | 3388 (94%) | 239 (7.9%) | 19  (1%) | 3109 (84.5%) | 301 (7.5%) | 100 (2.9%) |
|  | **Mid** | 174 (4.8%) | 2673 (88.2%) | 172 (9.1%) | 514 (14%) | 3639 (91%) | 685 (19.6%) |
|  | **Low** | 59 (1.6%) | 118 (3.9%) | 1699 (90%) | 56  (1.5%) | 57 (1.4%) | 2715 (77.6%) |
|  | **Total cells** | 3621 | 3030 | 1890 | 3621 | 3030 | 1890 |

*The result refers to the experiment presented in Fig. 2

** The result refers to the experiment presented in Fig S2A,B.

**Supplementary Table S1B: Number of active and inactive cells in each of the sorted sub-populations.**

|  | Sorted sub-populations  *MIT9313 old | | | Sorted sub-populations  **MIT9313 old | | |
| --- | --- | --- | --- | --- | --- | --- |
|  | **High** | **Mid** | **Low** | **High** | **Mid** | **Low** |
| % and number of active cells | 45  (68%) | 16  (22%) | 3  (3.1%) | 75  (64%) | 107  (51%) | 5  (6%) |
| % and number of inactive cells | 21  (31.8%) | 57  (78%) | 94  (97%) | 43  (36%) | 101  (49%) | 83  (94%) |
| Total cells | 66 | 73 | 97 | 118 | 208 | 88 |

*The result refers to the experiment presented in Fig. 2

** The result refers to the experiment presented in Fig S2A,B

**Supplementary Table 2:** **Coefficient of variation**

| **Strain and growth stage** | | **Sub-population** | **Illumination** | **Number of cells** | **C_cov_** | | **N_cov_** |
| --- | --- | --- | --- | --- | --- | --- | --- |
| **Batch culture** | |  |  |  | |  |  |
| MIT9313, exponential growth* | | High | Constant light  27 μE | 158 | | 0.92 | 0.89 |
| MIT9313,  old culture** | | High |  | 66 | | 1.08-1.29 | 0.92-0.87 |
|  |  | Mid |  | 73 | | 2.19-2.37 | 1.63-1.09 |
|  |  | Low |  | 97 | | 3.77-7.11 | 1.83-0.85 |
| MIT9313,  old culture*** | | High | Photo-period  12:12 L/D, 27 μE | 86 | | 2.65 | 1.02 |
|  |  | Mid |  | 171 | | 5.15 | 0.99 |
|  |  | Low |  | 73 | | 0.69 | 2.33 |
| Med4  Exponential growth**** | 3h |  | Constant light 27 μE | 489 | 0.40 | | 0.25 |
|  | 6h |  |  | 189 | 0.31 | | 0.25 |
|  | 12h |  |  | 780 | 0.39 | | 0.38 |
|  | 24h |  |  | 77 | 0.32 | | 0.28 |
| **Killed cells** | |  |  | 114 | | 2.63 | 0.09 |

*The results for MIT9313 exponential growth refer to the experiment presented in Fig. S3.

**The results for old MIT9313 cultures under constant light refer to the two experiments presented in Fig. 2 and Fig. S2A, B.

*** The results for Old MIT9313 cultures under L/D refer to the experiment presented in Fig. S2C-F.

**** The results for MED4 cultures refer to the experiment presented in Fig. S7.

For comparison, COVs of ~0.5-0.6 were recorded for uptake of different 15N-labeled N sources by the dinoflagellate *Prorocentrum minimum* *(6)*. Significantly larger variation was observed in field populations of photosynthetic picoeukaryotes, *Synechococcus* and *Prochlorococcus*, with *Prochlorococcus* revealing a COV of ~0.2-0.4 for ^13^CO_2_ and ~0.1-0.7 for ^15^NH_4_ (7).

**Supplementary figures**

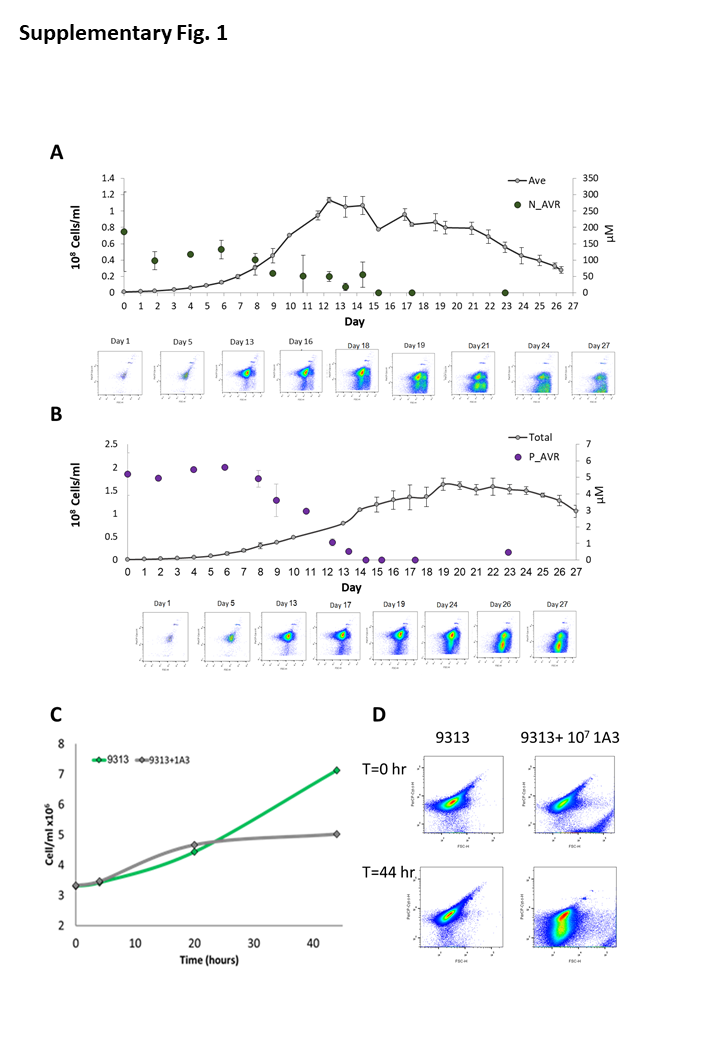

**Figure S1:** Sub-populations of low-chl cells emerge in *Prochlorococcus* cultures under N and P starvation, and when inhibited by heterotrophic bacteria. (A, B) Batch cultures of strain MIT9312 grown under conditions where stationary phase is induced by N and P starvation (panels A and B, respectively). The data shown are from (8, 9). (C) Batch cultures of *Prochlorococcus* MIT9313 grown axenically and in co-culture with a heterotrophic bacterium, *Alteromonas* HOT1A3. (D) FCM of *Prochlorococcus* populations at two time points (‘0’ and ‘44’ hours). Data are from Aharonovich and Sher (10).

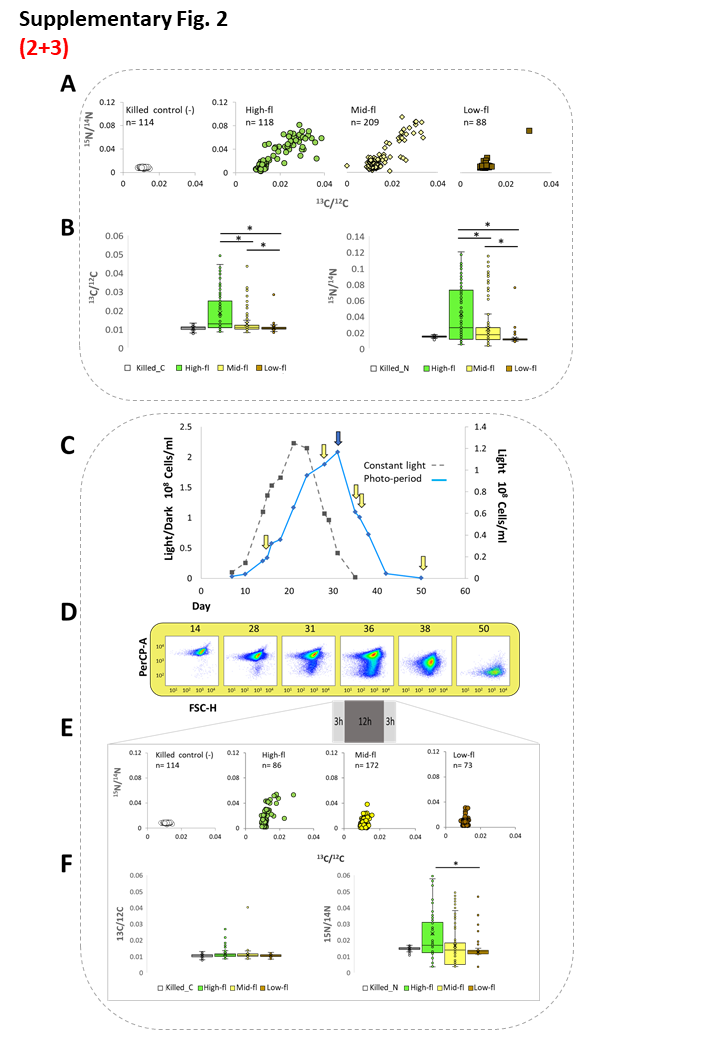

**Figure S2.** Independent experiments showing the metabolic activity of sorted sub-populations under constant light and a day/night cycle. (A, B) An independent experiment confirming differences in the metabolic activity of sorted sub-populations of MIT9313 by NanoSIMS. *Prochlorococcus* MIT9313 cultures were grown for 44 days in Pro99 and labeled with H^13^CO_3_^-^ and ^15^NH_4_^+^ for 18h. (A) Scatterplot of ^13^C/^12^C and ^15^N/^14^N ratios obtained from NanoSIMS analysis of each subpopulation (indicating the number of events detected from multiple fields). (B) Boxplot represent the variances of ^13^C/^12^C and ^15^N/^14^N in each sub-population. The three populations were statistically different (Kruskal-Wallis test, p<0.001, asterisks show significant differences in comparisons between each of the two populations using the Mann-Whitney U test, p<0.001). (C-F). NanoSIMS analysis for metabolic activity of *Prochlorococcus* chlorotic sub-populations under light/dark growth conditions. For *Prochlorococcus*, cell physiology is strongly entrained by the diel cycle, with cells typically dividing early during the night (11). As the cultures grown for these experiments presented in Fig 2 and Fig S2A and B were grown under conditions of constant light, it was possible that heterogeneity in C and N uptake rates was due to the presence of cells at different cell cycle stages. To test this hypothesis, we repeated this experiment using an MIT9313 culture grown under 12:12 light/dark conditions. (C) MIT9313 growth curve under 12 h light and 12 h dark (The arrows mark the days shown in panel D.) The dashed line shows the growth under constant light, for comparison. (D) A time series of flow cytometry scattergrams from the tested MIT9313 culture. The x-axis is Forward Scatter (FSC, a proxy for cell size), the y-axis is the chlorophyll autofluorescence of the cells. The appearance of chlorotic sub-population observed from the late exponential phase (Day 31). (E) Scatterplot of ^13^C/^12^C and ^15^N/^14^N ratios obtained from NanoSIMS analysis following 18 h incubation 3h L/12h D/3h L on day 36. (F) Boxplot f ^13^C/^12^C and ^15^N/^14^N enrichment in each subpopulation. Glutaraldehyde killed cells used as a negative control. Lines represent the median, X represents the mean, box borders are 1^st^ quartiles and whiskers represent the full range. The three populations were statistically different for N uptake (Kruskal-Wallis test, p<0.001) but not for C uptake (p=0.06). Significant differences between each two populations (Mann-Whitney U test) are shown (* - p<0.001). *The C uptake rate, as well as the cell-cell heterogeneity, were lower under these conditions, potentially because the cells were in darkness for 12 out of the 18 hours of labeling, including the “evening” and “morning” periods when the cell’s photosynthetic machinery is not running at its maximal capacity (11). In contrast, the N uptake rate remained high under the light-dark cycle. This suggests that NH_4_ uptake in Prochlorococcus is decoupled from photosynthesis and occurs during both light and dark periods, unlike amino acid uptake which occurs in Prochlorococcus primarily during the day (12).*

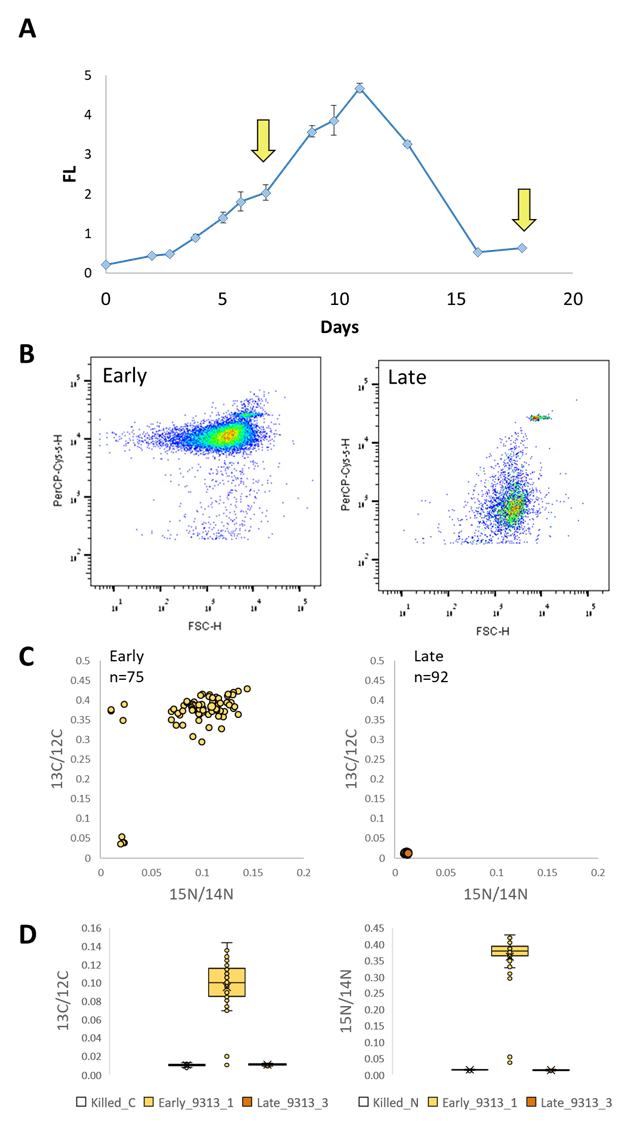

**Figure S3.** NanoSIMS analysis of *Prochlorococcus* MIT9313 from exponentially-growing, nutrient-replete cultures compared to late, post-decline stage in batch culture. We measured the N and C uptake rates in MIT9313 cultures for which the stationary stage is induced by N starvation (low-N Pro99, N:P ratio = 2, (8)). (A) Growth curve monitored via the chlorophyll auto-fluorescence. Arrows indicate the heavy-nutrient labeling time points (days 7 and 18). (B) FCM scatterplots of populations of 24hr post labelling. (C) Scatterplot of ^13^C/^12^C and ^15^N/^14^N ratios obtained from NanoSIMS analysis, representing single cell uptake. (D) Boxplot f ^13^C/^12^C and ^15^N/^14^N enrichment in each population, compared to killed cells in the control.

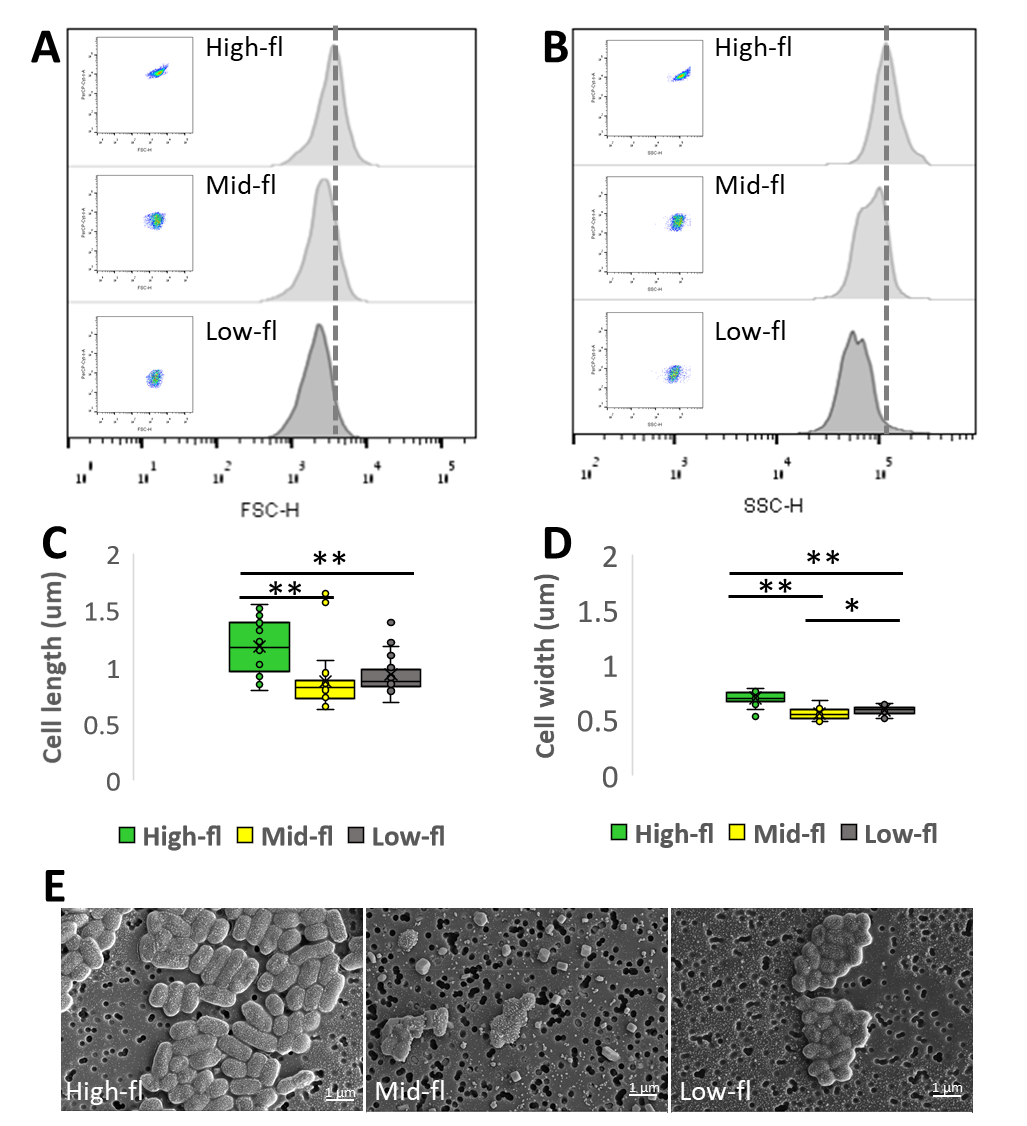

**Figure S4:** Sorted cells belonging to different sub-populations of *Prochlorococcus* MIT9313 vary in size. A late-exponential phase MIT9313 culture was fixed using glutaraldehyde, analyzed using flow cytometry, and the three different sub-populations sorted and observed by SEM. (A, B) Histograms representing changes in cell size (Forward scatter, FSC, panel A) and complexity (Side Scatter, SSC, panel B) as measured by flow cytometry. (C, D) Boxplots of variations in cell length (C) and cell width (D) as measured from SEM images of sorted populations. High-fl (n=27), Mid-fl (n=23), Low-fl (n=24). The three subpopulations were statistically different (Kruskal-Wallis test, p<0.001). Significant differences between each of the two populations (Mann-Whitney U test) are shown (** - p<0.001, * - p<0.05). Lengths values from Mid and Low were not significantly different. (E) SEM image of sorted sub-populations, scale bar is 1um. The small square objects in the middle panel are salt crystals.

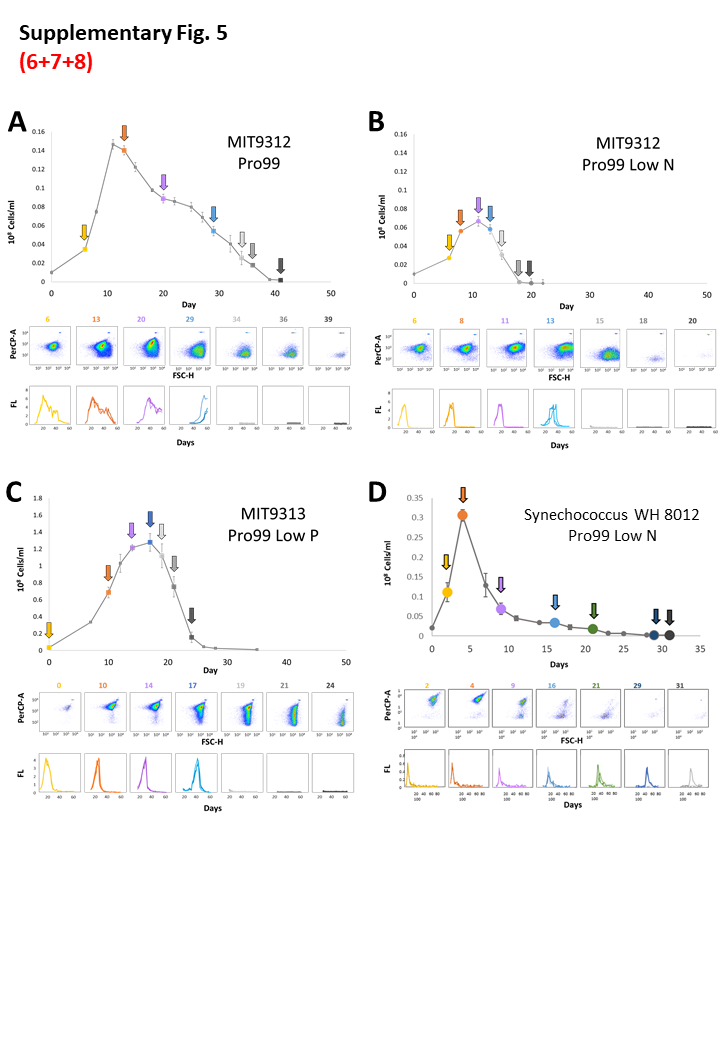

**Figure S5:** Time-dependent changes in viability of *Prochlorococcus* and *Synechococcus* cells from different strains transferred into fresh media at different life cycle stages and different media. In each panel, a growth curve is shown above flow cytometry scatterplots of specific time points (marked by arrows on the growth curves) and growth curves of cells being transferred at different times to new, nutrient-replete media (assessed via bulk culture fluorescence). Error bars on the top curves (cell numbers) represent mean and standard deviation of biological triplicates. (A, B) Time-dependent changes in viability of *Prochlorococcus* MIT9312 cells transferred into fresh media at different life cycle stages in Pro99 (A) and when stationary stage is induced by N starvation (B). The experiment in panel A was performed in Pro99 media commonly used for *Prochlorococcus* culturing (A) and in panel B under conditions where stationary stage is induced by nitrogen starvation (2:1 N/P ratio in the growth media, (8)). Note that, similar to strain MIT9313 (Fig 3), under nitrogen starvation, the cultures shift rapidly from being comprised primarily of high-fl cells (day 13 in panel B, early stationary phase) to mainly mid-fl cells, with essentially no high-fl cells (day 15 in panel B). Cells could not re-grow when transferred after more than 29 days in Pro99 and 13 days of nitrogen starvation. This suggests that, in this strain, low-fl cells are non-viable. (C) A batch culture of *Prochlorococcus* MIT9313 where stationary stage is induced by P starvation. The media used is Pro99 where P concentrations have been reduced 8-fold, leading to an N/P ratio of 144:1. Cells could not re-grow when transferred after more than 17 days. High-fl cells were still seen on days 19 and 21, when the cultures could not be transferred, suggesting that the high-fl cells are not necessarily viable. (D) A marine *Synechococcus*, strain WH 8012, survives much longer under N starvation than *Prochlorococcus*. This experiment was performed in Pro99 media in which N concentrations have been reduced 8-fold, leading to an N/P ratio of 2:1, similar to panel B, above (8). Under these conditions, chlorosis occurs faster than in Pro99, and thus in this experiment high-fl and low-fl were not observed together (compare with Fig 1C, where the same strain was cultured in Pro99). 2/3 cultures survived transfer on day 29 and 1/3 survived on day 31. Thus, in contrast to all tested *Prochlorococcus* strains, *Synechococcus* WH 8102 could be transferred long after entry into stationary phase, when only low-fl cells were observed. This suggests a fundamentally different relationship between cell chlorophyll fluorescence and viability (the ability to survive culture transfer in this strain).

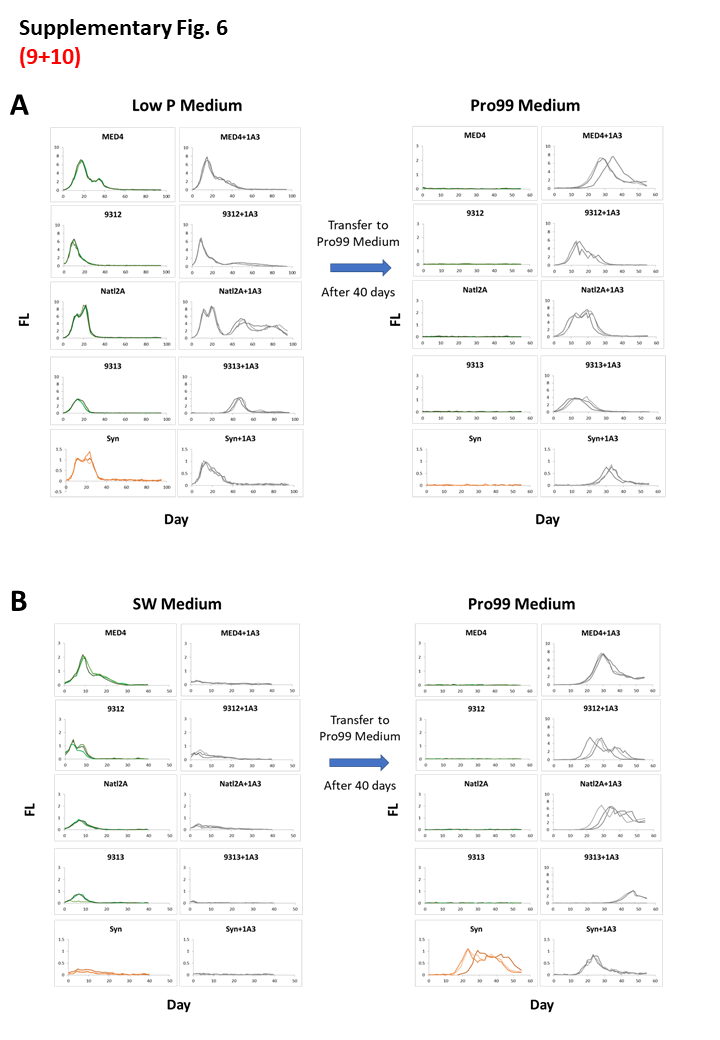

**Figure S6:** Co-culture with a heterotrophic bacterium, *Alteromonas* HOT1A3, enables multiple *Prochlorococcus* strains to survive long-term P starvation (A) and when cultured in natural seawater (B). Each panel shows the results of two consecutive experiments. The first experiment is shown on the left side, where either axenic cultures (left column, in green or orange) or co-cultures grown with *Alteromonas* HOT1A3 (right column, grey lines) are inoculated into the tested media. After 40 days, 1ml from each culture was transferred into fresh Pro99 media (right side of each panel), and the new cultures were monitored for an additional 50 days. In these plots, each line shows a replicate culture. (A) Culture survival under P starvation. PO_4_ concentration in the media was set to 6.25μM, leading to an N:P ratio of 128 rather than 16). Note the differences in the units on the y-axes. The length of starvation is longer in this experiment compared with the one presented in Supplementary Figure 5D (where *Synechococcus* WH 8102 cultures grew when transferred into new media for up to 31 days in N-starved cultures). (B) Culture survival in natural seawater (filtered but with no nutrients or trace metals added). Growth under these conditions is likely due to nutrient carryover from the original cultures, which is less than 40μM NH_4_ and 2.5μM PO_4_ (cultures were not centrifuged and resuspended in seawater to avoid cell stress). Note that, under these conditions, growth of *Prochlorococcus* in co-culture was reduced, potentially due to competition for inorganic nutrients by *Alteromonas*.

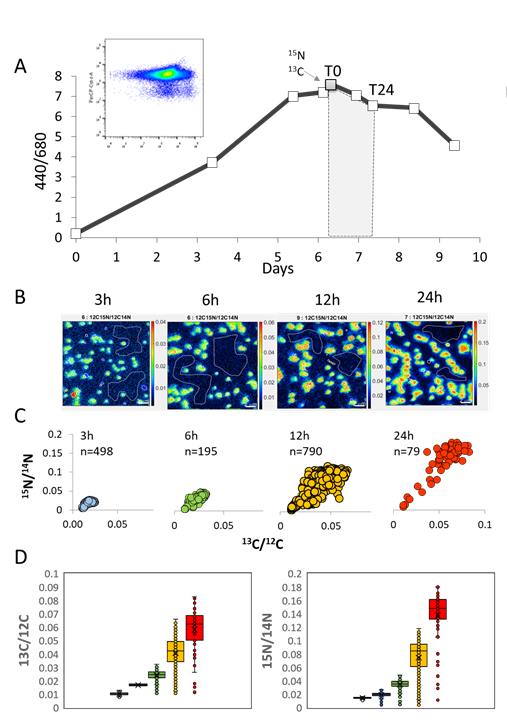

**Figure S7:** Optimization for measuring metabolic activity of *Prochlorococcus* by NanoSIMS (A) Growth curve of *Prochlorococcus* MED4 measured by fluorescence 440/680nm. Labeled nutrients (H^13^CO_3_^-^ and ^15^NH_4_^+^) were added after day 6 and sampled for NanoSIMS at 3h, 6h, 12h, and 24h. Insert shows the cell population at the time of labeling (T0) by FCM. (B) NanoSIMS images of ^15^N/^12^C analysis of cell for measuring metabolic activity. (C) Scatterplot of ^13^C/^12^C and ^15^N/^14^N ratios obtained from NanoSIMS analysis at each time point (indicating the number of events detected from multiple fields). (D) Boxplots represent the variances of ^13^C/^12^C and ^15^N/^14^N at each time point.
